# Geographic and seasonal variation of the *for* gene reveal signatures of local adaptation in *Drosophila melanogaster*

**DOI:** 10.1101/2023.02.19.529142

**Authors:** Dylan J. Padilla Perez

**Affiliations:** School of Life Sciences, Arizona State University, Tempe, Arizona 85287, USA

**Keywords:** foraging mode, landscape genetics, polymorphism, heterogeneous environments, demographic history

## Abstract

In the early 1980s, the observation that *Drosophila melanogaster* larvae differed in their foraging behavior laid the foundation for the work that would later lead to the discovery of the foraging gene (*for*) and its associated foraging phenotypes, rover and sitter. Since then, the molecular characterization of the *for* gene and our understanding of the mechanisms that maintain its phenotypic variants in the laboratory have progressed enormously. However, the significance and dynamics of such variation are yet to be investigated in nature. With the advent of next-generation sequencing, it is now possible to identify loci underlying adaptation of populations in response to environmental variation. Here, I present results of a genotype-environment association analysis that quantifies variation at the *for* gene among samples of *D. melanogaster* structured across space and time. These samples consist of published genomes of adult flies collected worldwide, and at least twice per site of collection (during spring and fall). Both an analysis of genetic differentiation based on *F_st_* values, and an analysis of population structure revealed an east-west gradient in allele frequency. This gradient may be the result of spatially varying selection driven by the seasonality of precipitation. These results support the hypothesis that different patterns of gene flow as expected under models of isolation by distance and potentially isolation by environment are driving genetic differentiation among populations. Overall, this study is essential for understanding the mechanisms underlying the evolution of foraging behavior in *D. melanogaster*.

## Introduction

Species distributed over broad geographic ranges face drastic variation in environmental conditions. Such species provide an opportunity to understand the mechanisms underlying evolutionary responses to environmental variation. *Drosophila melanogaster* represents a good example of a species with a widespread distribution range, being found on every continent and most islands (Markow and O’Grady, 2005). Since the early 1900s until the present days, this species has been a central model to understand the genetic basis of adaptation (Flatt, 2020). Identifying genes underlying adaptation of populations is a key issue in evolutionary biology (Tenaillon and Tiffin, 2008). For instance, the foraging gene (*for*) is best known for underlying different foraging strategies adopted by *D. melanogaster* in response to environmental variation (Anreiter and Sokolowski, 2019). Detailed characterization of this gene in laboratory experiments revealed the existence of two phenotypic variants: rover and sitter (Sokolowski, 1980). In the presence of nutritive yeast, larval rovers travel longer distances and eat less than sitters do. This difference is not evident in the presence of a non-nutritive agar medium (Kaun et al., 2007), suggesting a gene-by-environment interaction (Burns et al., 2012). Similarly, Pereira and Sokolowski (1993) showed that adult rovers walked significantly farther from the food source after eating than did sitters, but there were no differences in walking behavior in the absence of food. This behavioral variation should also be observed in the wild, because food availability varies temporally and spatially across the range of *D. melanogaster*. Because these behavioral differences between rovers and sitters arise from genetic variation at the *for* gene, one should expect the frequency of alleles of this gene to vary over space and time.

Extensive geographic and temporal variation in various genetic markers has been identified in *D. melanogaster* (Hoffmann and Weeks, 2007; Kapun et al., 2020; Sezgin et al., 2004). In some cases, clinal genetic variants have been directly linked to phenotypic variants (Lee et al., 2013; Paaby et al., 2014, 2010; Schmidt et al., 2008). For example, associations between the *Adh* allozyme locus and development, as well as the *hsp70* gene and heat knockdown resistance have been discovered from a mid-latitude population along the east coast of Australia. There is also good evidence indicating that *D. melanogaster* evolved Bergmann’s clines on more than one continent. When raised in a common environment, both South American and Australian flies from high latitudes developed faster and matured at larger sizes than their counterparts from low latitudes (James et al., 1997). Similarly, seasonal stresses at high latitudes, such as overwinter survival and ephemeral food resources, impose strong selection for increased body size and somatic maintenance, whereas warm climates with ample feeding opportunities favor reduced size and decreased maintenance (James et al., 1997; Paaby and Schmidt, 2009). Because the foraging behavior of organisms influences their ability to gather and assimilate food to fuel their somatic maintenance, one may expect the allele frequency of the *for* gene to differ in genotypes living at high latitudes and over stressful time (e.g., winter), relative to those living at lower latitudes and less stressful conditions.

Although recent studies have provided good insights on the maintenance of the *for* gene polymorphism in the laboratory, the significance and dynamics of such variation are yet to be investigated in nature. Studies based on experimental evolution suggest that density-dependent selection during the larval stage of Drosophila, rather than the adult stage, can maintain genetic variation in foraging behavior. For instance, a high larval density favors the rover variant and low larval density favors the sitter variant (Sokolowski et al., 1997). Similarly, negative frequency-dependent selection under low food condition promotes the increase in frequency of one variant when it is rare relative to the other variant (Fitzpatrick et al., 2007). In addition, Anreiter et al. (2017) showed evidence of an epigenetic regulator of the *for* gene (the *G9a* methyltransferase) responsible for rover-sitter differences in adult foraging behavior. Specifically, they showed that rovers bearing *G9a* alleles, had significantly greater foraging success than sitters, but this rover–sitter difference disappeared when they carried *G9a* null alleles. In sum, researchers have now detailed advances on the mechanisms that produce variation in the Drosophila *for* gene under controlled conditions. Yet, our understanding of the environmental drivers of the genetic variation that influences the foraging behavior of flies in nature remains unknown. A good approach to fill this gap is referred to as genotype-environment association analysis (Joost et al., 2007; Rellstab et al., 2015), a statistical tool used in the fields of landscape genetics and population genomics to identify the environmental factors that have shaped present-day genetic variation, and the gene variants that drive local adaptation.

Here, I present results of the first genotype-environment association analysis that quantifies variation at the *for* gene among samples of *D. melanogaster* structured across space and time. These samples consist of published genomes of adult flies collected worldwide, and at least twice per site of collection (during spring and fall). Thus, not only could I test whether the allele frequency of the *for* gene changes geographically, but also in response to seasonality. In natural populations, quantitative traits that exhibit continuous geographic variation are often associated with specific ecological variables reflecting selective pressures acting on individual phenotypes. Though, evidence for local adaptation to continuous environments can be efficiently detected if there is highly significant association with the environmental variables at some loci compared with the background genomic variation (Kelley et al., 2006). Accordingly, a screening procedure to detect locus-specific signatures of positive selection that accounts for chromosome-wide patterns of polymorphism should then provide robust insights. This study provides an analytical framework that compares the allelic variation at the *for* gene relative to the variation of the whole chromosome 2L. In doing so, I found some degree of genetic differentiation among samples within America and Europe based on a genetic admixture analysis, suggesting a total of 9 ancestral populations (*k* = 9) with unique patterns of allelic variation at the *for* gene. Interestingly, I found a lower degree of genetic differentiation among populations through the screening of the whole chromosome 2L (*k* = 8). The results suggested that seasonality strongly contributed to the genetic differentiation of populations, but its effect depends on the site of collection.

## Materials and Methods

### Data source

The data used in this study represent a subset of a dataset assembled by the Drosophila Genome Nexus project (DGN), the European Drosophila Population Genomics Consortium (DrosEU), and the Real Time Evolution Consortium (DrosRTEC). This dataset is known as the Drosophila Evolution over Space and Time (DEST), and it is coupled with environmental metadata (see https://github.com/DEST-bio/DEST_freeze1/blob/main/populationInfo/samps_10Nov2020.csv). It constitutes the most comprehensive spatiotemporal sampling of *D. melanogaster* populations to date, including estimates of genome-wide allele frequencies for more than 270 populations. Collectively, the data are based on sequencing of more than 13,000 flies collected around the world over multiple years (Kapun et al., 2021). Importantly, the analyses of this study relied on pooled samples stored in the Genomic Data Structure (GDS) file named “dest.all.PoolSNP.001.50.10Nov2020.ann.gds”, available at https://dest.bio/data-files/SNP-tables. Pooled samples from this file consist of 33-40 wildcaught males, which are more easily distinguishable morphologically from similar species than females are. Detailed information on quality control, mapping, SNP calling and dataset merging can be obtained from the DEST dataset mapping pipeline (see https://dest.bio/sourcecode).

I filtered the GDS object to restrict my focus on Single Nucleotide Polymorphism variants (SNPs) at the *for* gene. I also performed a screening of the entire chromosome 2L to compare the levels of variation at the *for* gene relative to the variation of the genomic background. As annotated at https://flybase.org/ (genome assembly version 6.32), the *for* gene is found on the left arm of chromosome 2 from base pairs 3,622,074-3,656,953. I further filtered the reads to get a subset of pools collected at least once during spring and winter across multiple years, ranging from 2014 to 2019 (see Table S1 for the exact number of season per locality). Although the data enabled me to assess the effect of seasonality on allelic variation at the *for* gene, I failed to test for an interaction between year and season, because both seasons could not always be sampled in the same year. Because the filtering process drastically reduced the number of pools in a few populations, I grouped together pools from the nearest neighboring localities such that the smallest populations had 5 pools. The resulting number of pools are distributed among 15 populations collected in Austria EU (*n* = 6), Charlottesville USA (*n* = 6), Denmark EU (*n* = 5), Esparto USA (*n* = 5), Finland EU (*n* = 6), France EU (*n* = 10), Germany EU (*n* = 12), Michigan-Wisconsin USA (*n* = 9), New York-Massachusetts USA (*n* = 6), Pennsylvania USA (*n* = 18), Russia EU (*n* = 6), Spain EU (*n* = 7), Tuolumne USA (*n* = 5), Turkey EU (*n* = 13), and Ukraine EU (*n* = 38). The DEST dataset provides the coordinates of each collection site, enabling one to obtain data about the seasonality of temperature and precipitation. These climatic variables were derived from the monthly values of temperature and rainfall for global land areas at a resolution of 30 s of a longitude/latitude degree spatial resolution (this is about 900 m at the equator) available at WorldClim v.2 (Fick and Hijmans, 2017). In addition, I extracted log-transformed values of net primary production (NPP)—the net amount of solar energy converted to plant organic matter through photosynthesis—measured in units of elemental carbon. This variable represents the primary source of trophic energy for the world’s ecosystems and can be obtained from the NASA Socioeconomic Data and Applications Center (Imhoff et al., 2004, sedac). These environmental variables were used in a genotype-environment association analysis as described below.

### Quality control

To avoid the influence of atypical observations driven by samples with low coverage, I applied a minimum effective coverage *N_eff_* filter of 28 (e.g., Nunez et al., 2022). I also removed pools and SNP loci with more than 20% missing genotypes. To do so, I used the *missingno* function from the “poppr” package of R v2.9.3 Kamvar et al. (2014). Subsequently, I tested for deviations from Hardy–Weinberg equilibrium (HWE) using the function *hw.test* from the “pegas” package of R v1.1 Paradis (2010). This function calculated p-values of an exact test with a Monte Carlo permutation of alleles (1,000 replicates). I considered loci to be out of HWE if they deviated significantly in more than 50% of populations. This decision was based on a “false discovery rate” correction of the p-values.

### Genetic differentiation and population structure

To test for genetic differentiation among populations at the level of the *for* gene, I estimated pairwise values of *F_st_* adopting the method suggested by Weir and Cockerham (1984). The *F_st_* values serve as a tool for describing the partitioning of genetic diversity within and among populations (Wright, 1931). For example, large populations among which there is much migration tend to show little differentiation with values of *F_st_* close to 0, whereas small populations among which there is little migration tend to be highly differentiated with values of *F_st_* close to 1. I used the function *gl.fst.pop* from the package “dartR” v2.7.2 (Gruber et al., 2018), running 1,000 bootstraps across all loci to generate 95% confidence intervals of the values of *F_st_*. When the confidence intervals did not include zero, the *F_st_* value in question was considered significantly different from zero.

I estimated population structure based on admixture proportions. I performed this procedure for both the *for* gene and the chromosome 2L. Specifically, the admixture analysis uses a sparse negative matrix factorization method (SNMF), available in the R package “LEA” v3.2.0 (Frichot and François, 2015). Similar to Bayesian clustering programs such as CLUSTER, the function *snmf* in LEA estimates individual admixture coefficients from a matrix of allele frequencies. Assuming *k* ancestral populations, the function provides least-squares estimates of ancestry proportions and estimates an entropy criterion to evaluate the quality of fit of the statistical model through cross-validation. The entropy criterion enables one to choose the number of ancestral populations that best explain the data. Specifically, I used the function *snmf* in LEA to compute individual ancestry coefficients for a number of ancestral populations (*k*), ranging between *k* = 1 and *k* = 10. For each value of *k*, I ran the algorithm 50 times and calculated the cross-entropy for each run.

### Seasonality analysis

To assess the effect of seasonality on allele frequencies at the *for* gene among populations, I used a constrained ordination analysis, which enables one to determine the genetic variation that can be explained by certain environmental variables or constraints. Accordingly, I performed a Constrained Correspondence Analysis (cca) through the function *cca* in the R package “vegan” v2.6-4. This function requires an independent variable (season), and a dependent variable (allele frequencies matrix). The output of this function decomposes the total variation into constrained and unconstrained components. To determine whether seasonality had a significant effect on the allele frequencies across populations, I used a permutation test using the *anova* method function. Accordingly, the data are permuted randomly and the model is refitted. When the constrained component in permutations is nearly always lower than the observed constrained component, one says that the constraint is significant.

### Detecting adaptation

To detect signatures of local adaptation, I performed a genotype-environment association analysis based on Latent Factor Mixed Models (LFMM) with a ridge penalty (Jumentier et al., 2022). Specifically, I first performed the analysis at the *for* gene alone, and then across the whole chromosome 2L, including *for*. The LFMM method is a univariate test that requires a matrix of allele frequencies as a dependent variable and an independent variable. Thus, I fitted three different models, each of which included a different predictor variable (e.g., precipitation, temperature, and NPP). The LFMM method also requires an estimate of the number of ancestral populations in the data (*k*), which relied on the admixture analysis discussed earlier. To correct for potential family-wise error rates, I fitted the models using the default genomic inflation factor correction (GIF) and a conservative False Discovery Rate (FDR) of 0.001 by converting the p-values to q-values. The FDR is the expected proportion of false positives among the list of positive tests (Storey and Tibshirani, 2003). For example, an FDR threshold of 0.001 applied to a dataset with an ideal p-value distribution would produce 0.1% false positives (false discoveries) among the set of positive tests. However, if the distribution of p-values deviates from the ideal distribution, this same FDR threshold of 0.001 would produce more or fewer false discoveries among the set of positive tests, depending on the skew of the distribution.

Finally, I explored patterns of isolation by environment (IBE) and isolation by distance (IBD). I performed these analyses only on European samples considering that isolation by distance results in differences between populations at opposite ends of a more or less continuous group of populations. Unlike the European samples, the American populations were narrowly distributed on the west and east coasts, with no populations spanning the center of the country. In such cases, the dispersal distances might be too large for isolation by distance to be accurately detected. The IBE and IBD analyses require the estimation of distance matrices such as a genetic distance matrix according to Nei (1987), and both a matrix with ecological distances (e.g., precipitation distance) and a matrix with geographic distances (coordinates) based on Euclidean distance. To do this, I used a Multiple Matrix Regression with Randomization analysis (MMRR), which incorporates multiple regressions and can be extended to any numbers of predictor variables (Wang, 2013). A multiple regression equation for distance matrices can be estimated using standard multiple regression technique, with the exception that tests of significance must be performed using a randomized permutation because of the non-independence of elements (Wang, 2013).

## Results

In general, none of the *for* loci nor populations were consistently out of Hardy-Weinberg equilibrium. I filtered out of the analyses only one sample that had more than 20% of missing genotypes, and 62 loci that had more than 20% of missing data. The quality control procedure generated a new dataset of 1,550 loci (SNPs), and 3,100 alleles from 151 pools distributed among 15 populations. Similarly, the quality control of the variants associated with the entire chromosome 2L resulted in a dataset of 931,161 loci, and 1,862,322 alleles from 151 pools distributed among 15 populations.

An analysis of genetic differentiation based on SNP variants of the *for* gene led to an overall *F_st_* value of 0.032, but the pairwise differences in *F_st_* ranged from 0 to 0.069. Interestingly, the analysis of genetic differentiation based on the *for f_st_* values revealed two major clusters separating the pools collected in America from those collected in Europe (Figure 1). However, significant differentiation was also observed within each continent.

**Figure 1:**
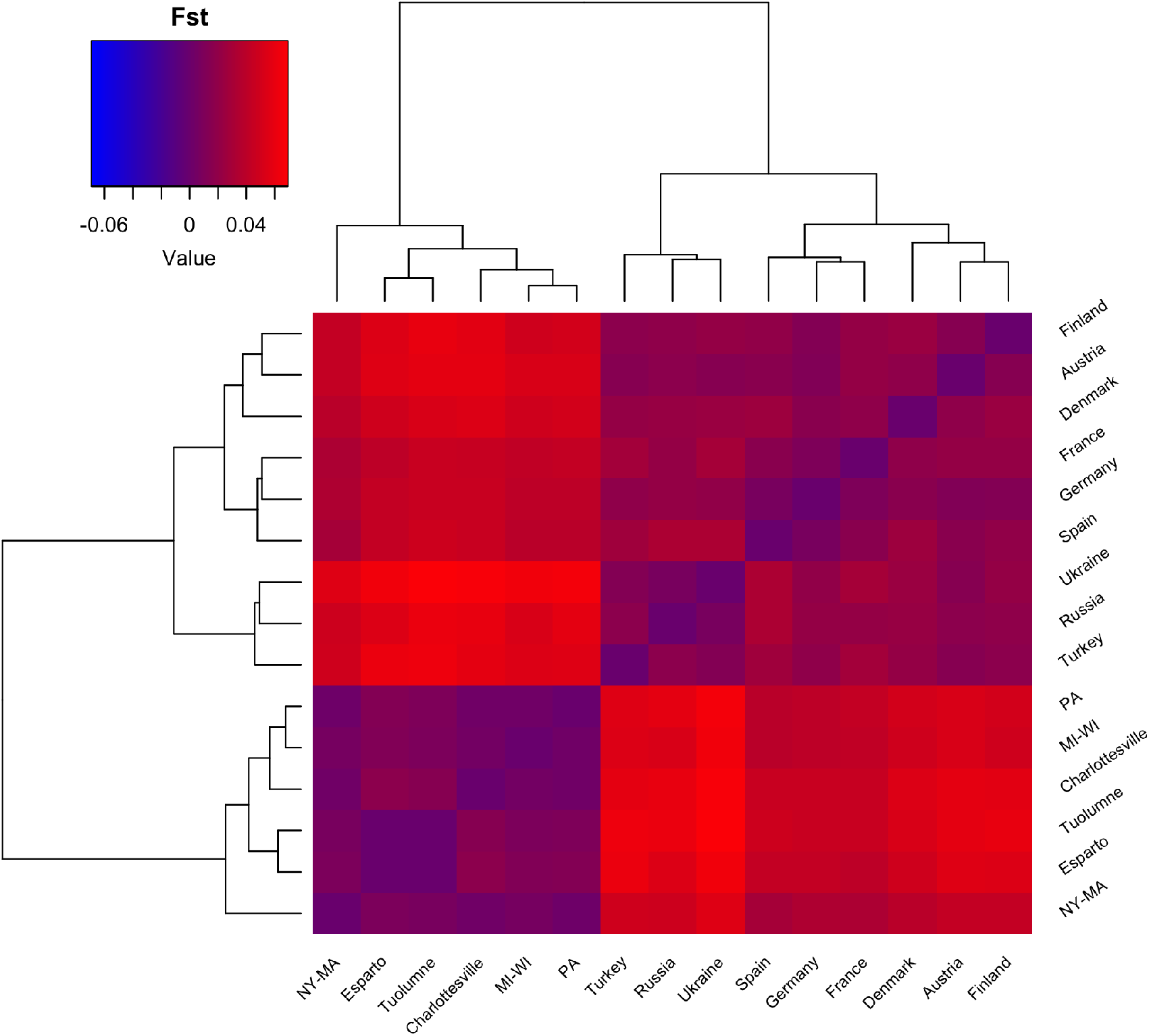
Heatmap depicting genetic differentiation among populations of *D. melanogaster* based on pairwise *F_st_* values. In some cases, samples from adjacent localities were combined to avoid low sample sizes. Abbreviations are as follows: PA = Pennsylvania; MI-WI = Michigan and Wisconsin; NY-MA = New York and Massachusetts.

The genetic admixture analysis suggested a pattern of genetic clusters that approximates that observed from the *F_st_* values. The *snmf* run at the *for* gene with *k* = 9 best accounted for population structure among all samples (Figure 2a). By contrast, a *snmf* run across chromosome 2L with *k* = 8 best accounted for population structure. Overall, I observed a distinct east-west gradient in population structure according to the admixture proportions (Figure 2b). This gradient seems to be stronger among the populations collected in America than those collected in Europe, although the structure of pools from Ukraine, Turkey, and Russia stands out (Figure 3).

**Figure 2:**
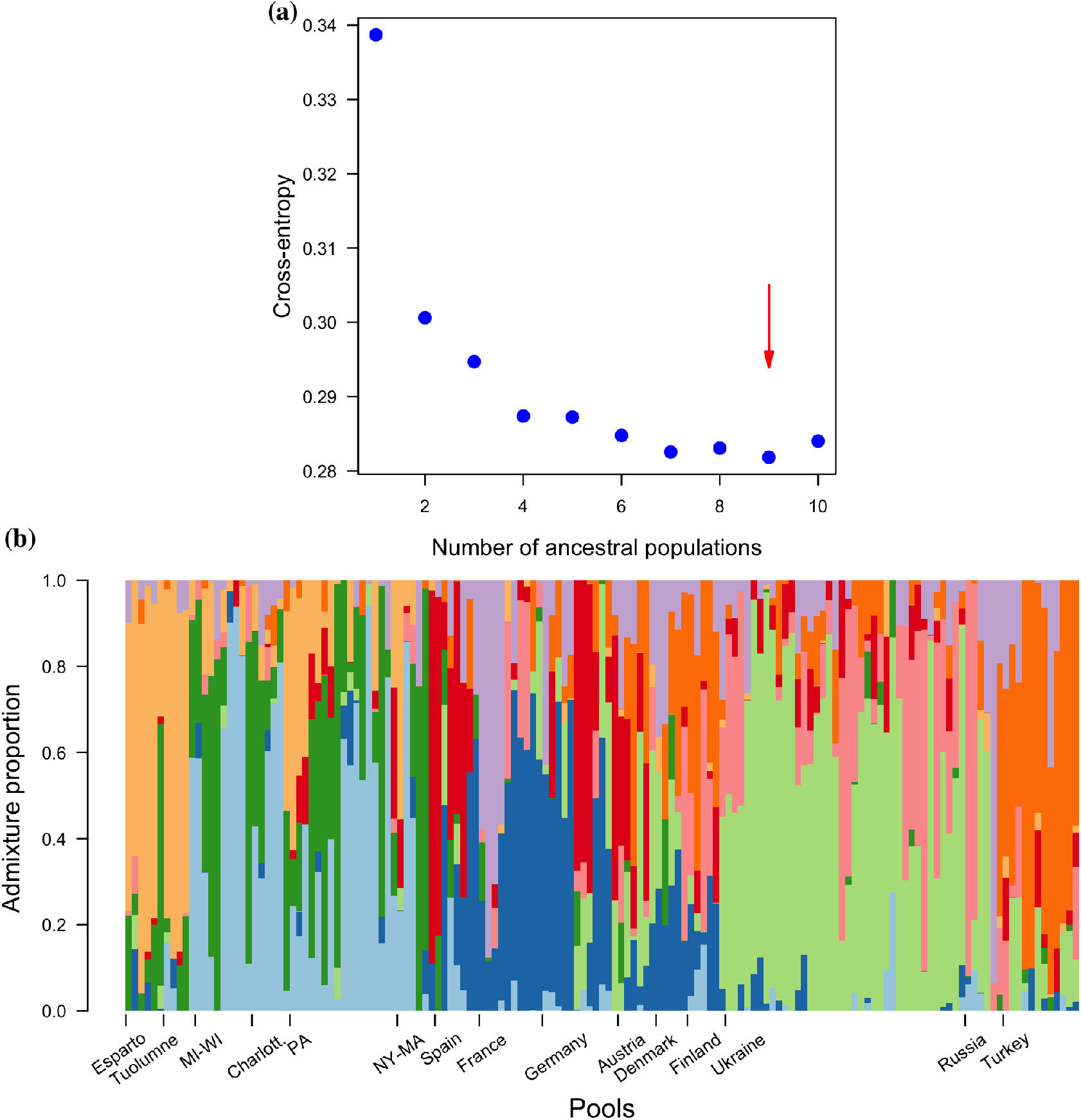
Population structure analysis based on individual ancestry coefficients for a number of ancestral populations. **(a)** Cross-entropy values for each *snmf* run with *k* ranging between *k* = 1 and *k* = 10. The red arrow indicates the most likely value of *k*. **(b)** Admixture proportion across populations of *D. melanogaster*. Colors indicate genetic clusters.

**Figure 3:**
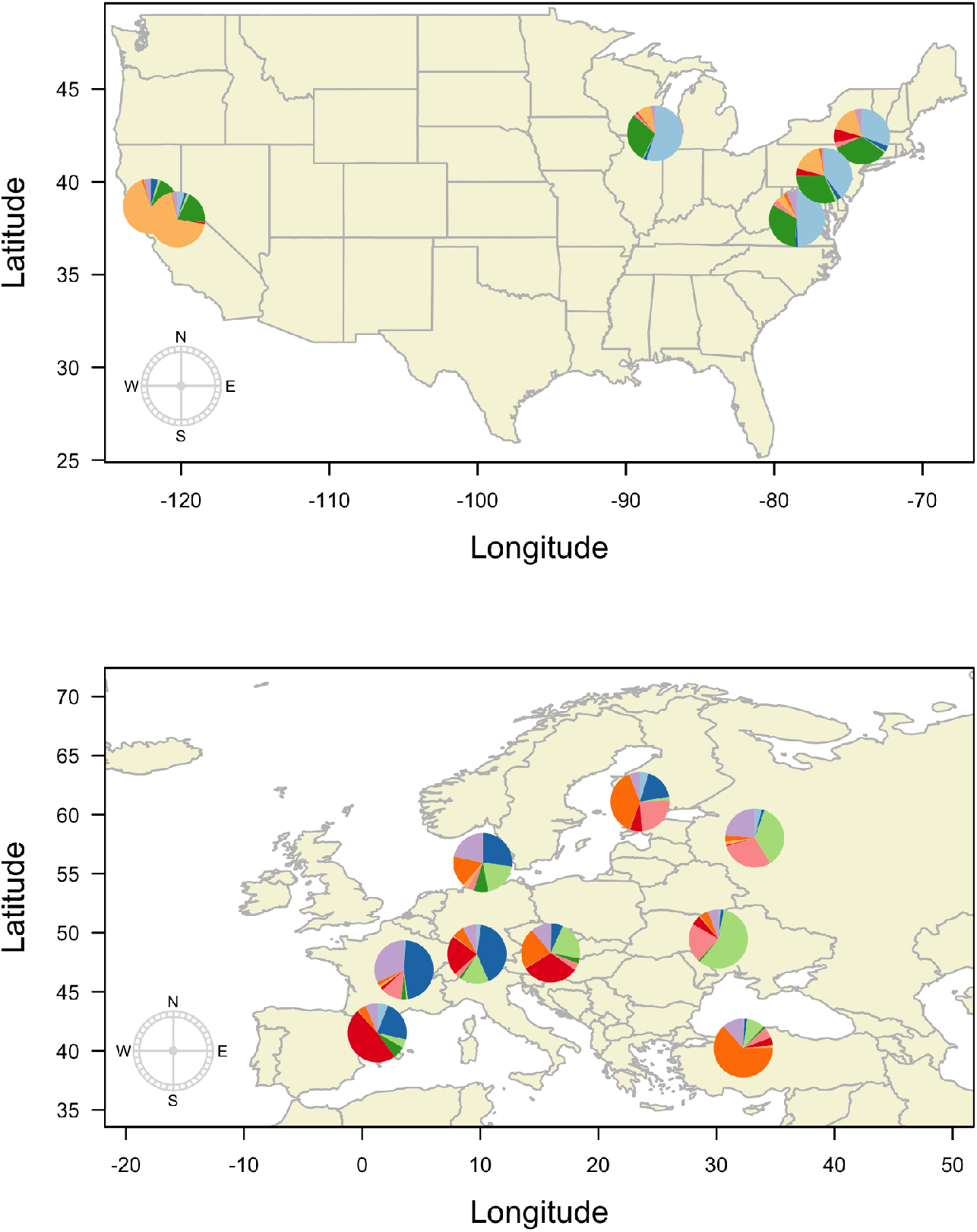
Mean admixture proportions across populations of *D. melanogaster* surveyed in America (upper panel) and Europe (lower panel). Colors in the pie charts represent genetic clusters.

According to the constrained correspondence analysis, an interaction between season and locality seemed to drive the difference in allele frequency across populations 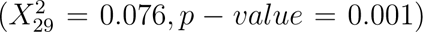. Despite the substantial divergence of populations in response to seasonality, some populations seem to be overlapping and cannot be completely separated by the effect of seasonality (Figure 4).

**Figure 4:**
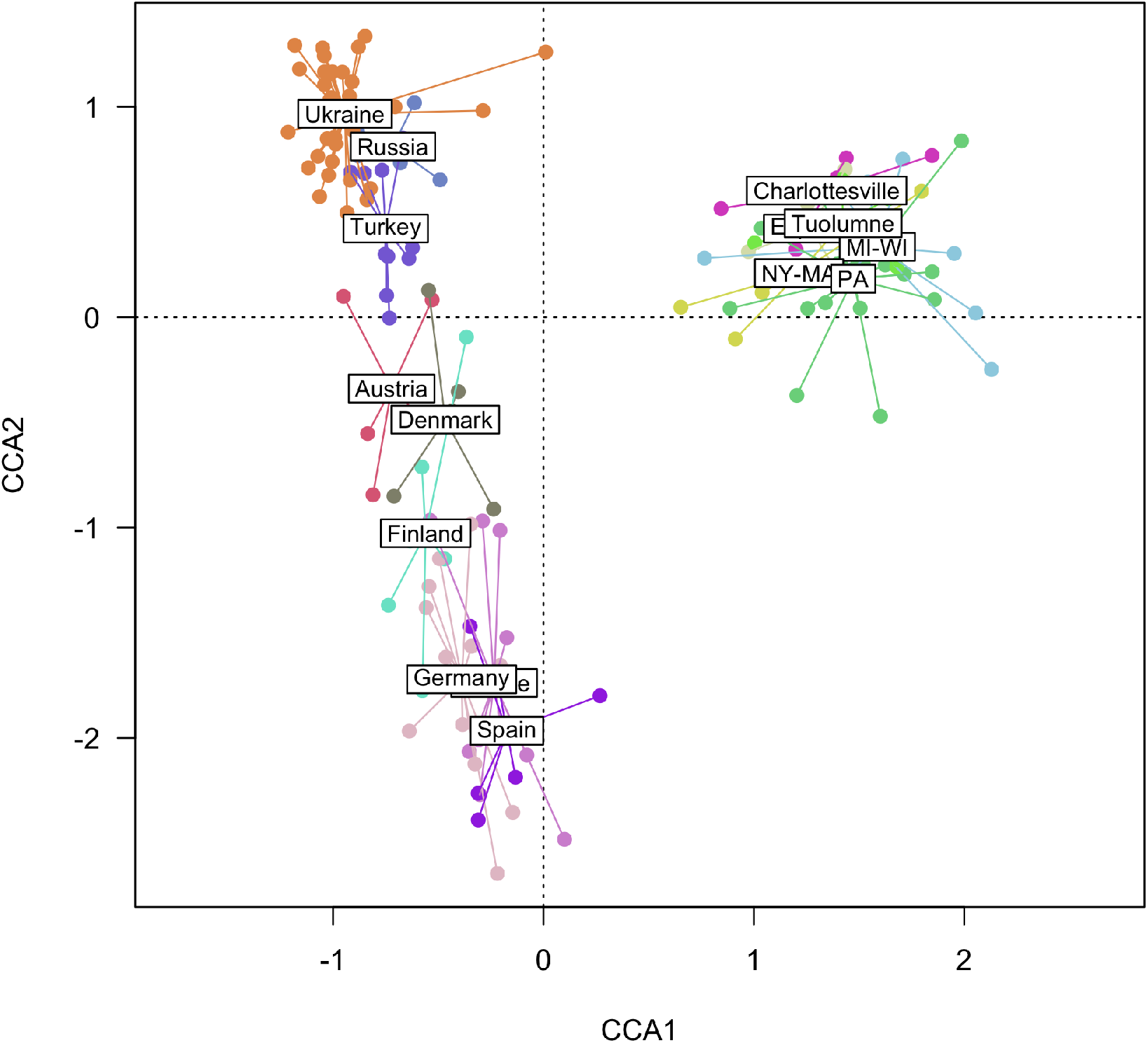
Constrained correspondence analysis depicting the difference in allele frequency across populations mediated by the interaction between season and locality. Filled dots represent pooled samples and “spider” diagrams connect the pools to the population centroid.

From the latent factor mixed models performed at the level of the *for* gene and the entire chromosome 2L, I extracted locus-specific z-scores and adjusted p-values (Figure 5). Using *k* = 9 as suggested by the admixture analysis at *for*, the default GIF correction, and an FDR threshold of 0.001, I detected 2 candidate SNPs under selection in response to precipitation (Figure 5a). Interestingly, I found no loci under selection based on the effects of temperature and NPP. By contrast, the screening of the whole chromosome 2L polymorphism using *k* = 8 revealed only 52 candidate SNPs under selection in response to precipitation. Importantly, one of the SNPs under selection previously discovered at the *for* gene was also observed through the analysis of the entire chromosome 2L (Figure 5b).

**Figure 5:**
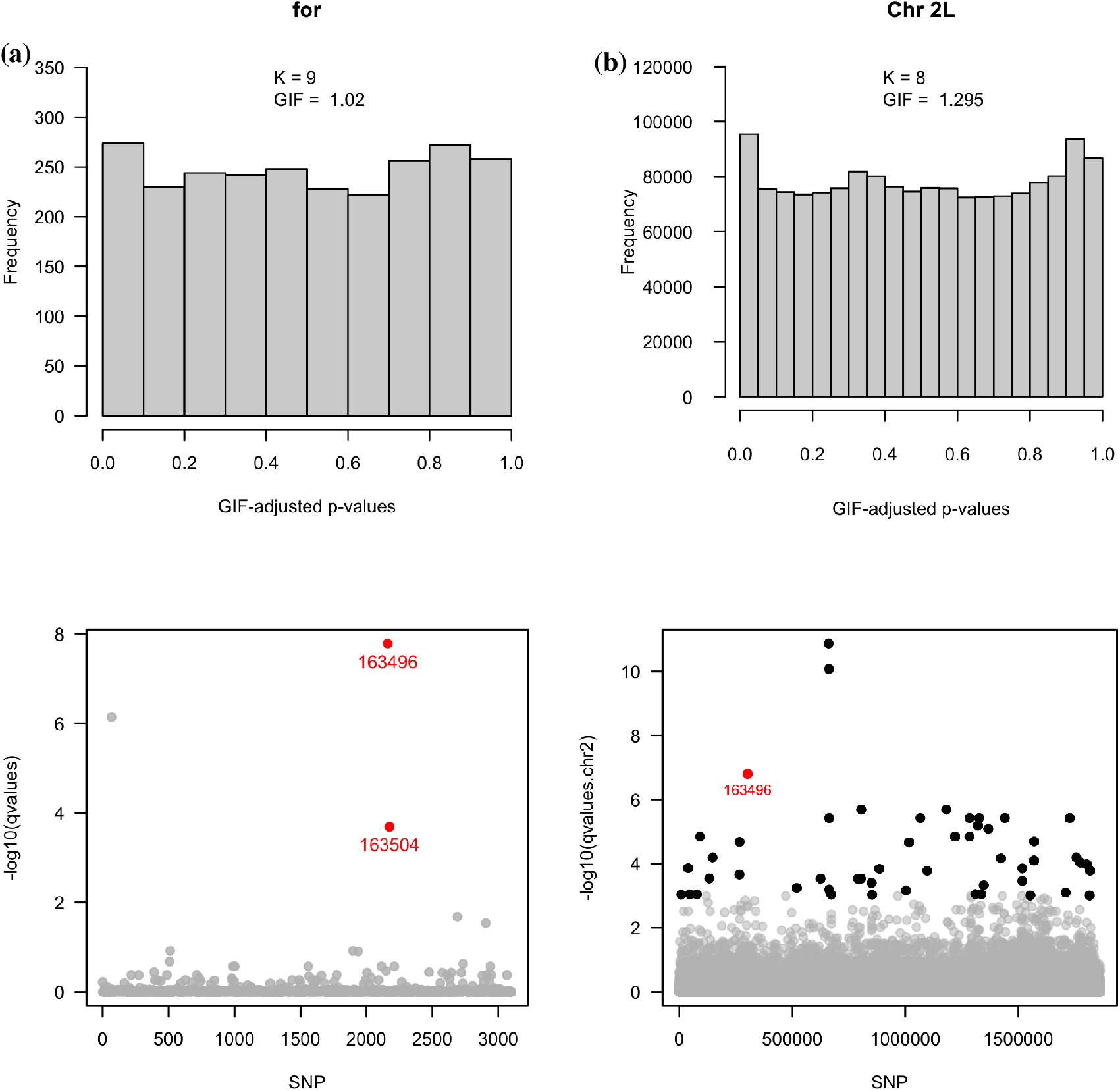
Genotype-environment association test based on a latent factor mixed model with ridge penalty. **(a)** Distribution of adjusted p-values using the default genomic inflation factor, and the *for* loci (SNPs) potentially affected by precipitation. Likewise, **(b)** displays the same analysis for the entire chromosome 2L. Bold dots represent loci considered to be under selection according to a False Discovery Rate of 0.001.

The IBE and IBD analyses based on the MMRR test showed an increase of genetic distance with geographic distance (*β* = 4.728 *×* 10*^−^*^3^, *t* = 5.430, *p − value* = 0.001). Similarly, the genetic distance increased with precipitation distance but the slope of the relationship was not significant (*β* = 1.568 *×* 10*^−^*^3^, *t* = 1.433, *p − value* = 0.185). These results suggested that populations are relatively similar at short distances, while this similarity decreases with geographic distance (Figure 6a-b).

**Figure 6:**
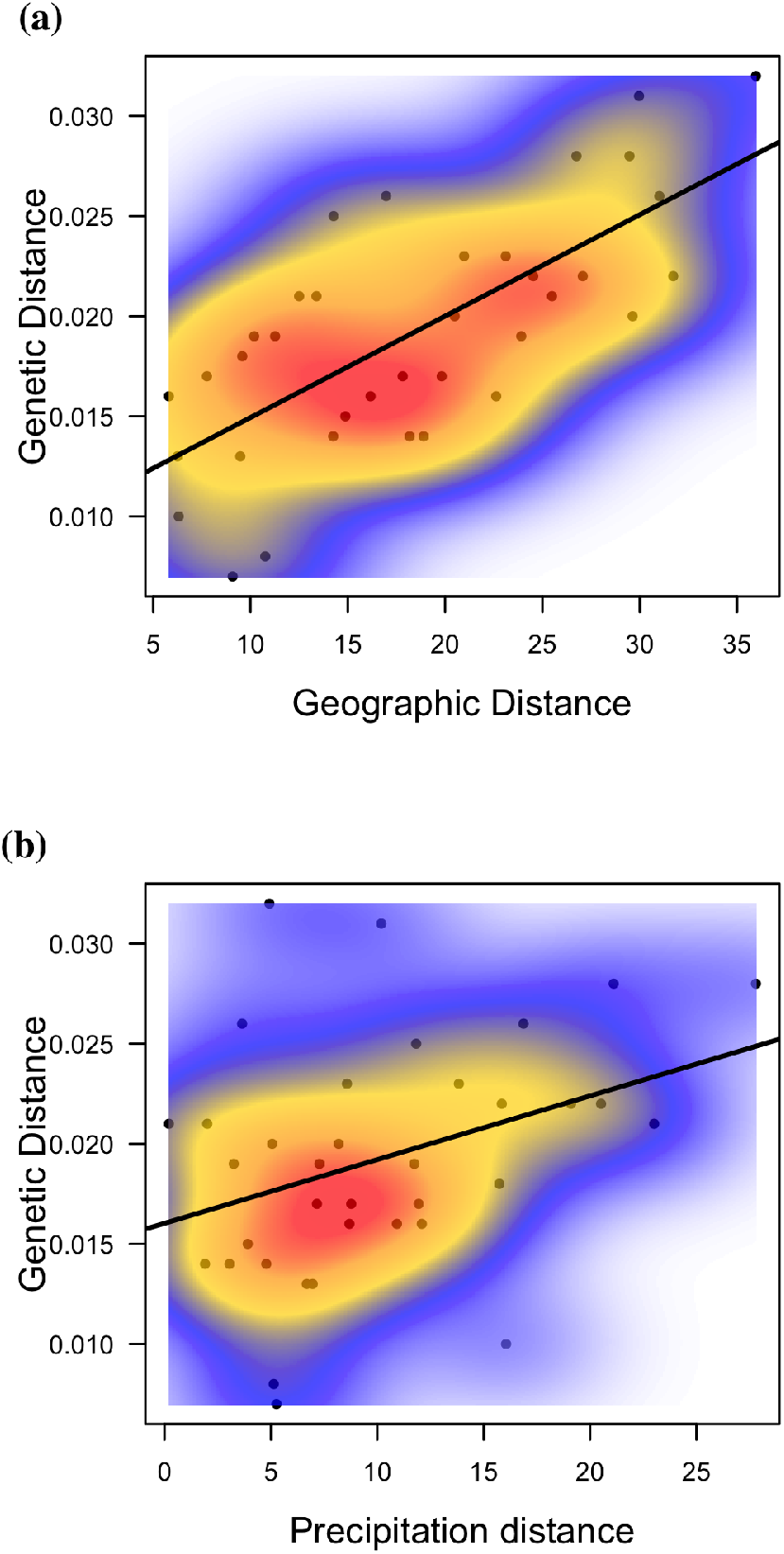
Patterns of isolation by distance and isolation by environment according to Multiple Matrix Regression with Randomization analysis (MMRR). **(a-b)** Geographic and precipitation distances as a function of genetic distance in European populations of *D. melanogaster*. Colors represent estimated probability densities.

## Discussion

This study shows that populations of *D. melanogaster* are geographically differentiated based on SNP variants of the foraging gene (*for*). Interestingly, the degree of genetic differentiation appears to be greater based on the allelic variation of the *for* gene relative to that of the genomic background. This finding was supported by estimates of the *F_st_* index and individual admixture proportions among populations. Patterns of genetic differentiation among populations might have been shaped by both neutral processes such as demographic history, and adaptive forces such as environmental heterogeneity. Some evidence indicate that North American populations of *D. melanogaster* have been affected by African and European migrations associated with human cargo transportation and fruit trade around the world, contributing to genetic admixture worldwide in recent years (Bergland et al., 2016; David and Capy, 1988; Kao et al., 2015; Lachaise et al., 1988). Numerous studies have also revealed clinal variation in phenotypes, chromosomal arrangements, and genotypes across environmental gradients for this species, implicating spatially varying selection (Flatt, 2016; Hoffmann and Weeks, 2007). This study suggests that genetic variation at the *for* gene seems to be influenced by seasonality, but this effect varies according to the site of collection. Although an effect of seasonality seems evident across populations, some populations cannot be separated by the effect of seasons. To understand such variation, the effect of natural selection should be disentangled from demographic events (Miller et al., 2020; Sexton et al., 2014; Smith et al., 2020), and other unobserved factors. Those unobserved factors include uneven sampling designs, genome-sequencing biases, relatedness among individuals, gene interactions that affect phenotypic variation, and linkage disequilibrium within haplotypes (François et al., 2016). To deal with these factors, I applied a stringent correction to the p-values across the statistical test performed in genotype-association analysis (Balding, 2006; Pearson and Manolio, 2008; Korte and Farlow, 2013). Displaying the empirical distribution of the adjusted p-values (see Figure 5) is a good way to show that confounding effects were removed from the analysis and that false discoveries could be controlled (Storey and Tibshirani, 2003; McCarthy et al., 2008). Once confounding errors are removed, adjusting for multiple comparisons could then be achieved through the application of FDR control algorithms (Benjamini and Hochberg, 1995; Storey and Tibshirani, 2003).

According to the genotype–environment association analysis, I found two potential intronic variants under selection at *for*. One of the *for* variants remained under selection when the allelic variation of the genomic background was accounted for. The genotype–environment association analysis identifies loci that show strong correlations with one or more environmental variables, and corrects for confounding effects when the environment is correlated with population structure. When modeling the effect of the environment on genetic variation at the *for* gene, I found that seasonality strongly influences the allelic differentiation across populations of *D. melanogaster*. But the effect of seasonality is actually mediated by precipitation and temperature. Generally, abundant precipitation and radiation increase the rate of photosynthesis (Cramer et al., 1999), which translate into higher NPP conditions required for growth and reproduction of organisms at higher trophic levels. Consistent with this idea, the genotype–environment association analysis revealed that precipitation strongly influenced the variation in allele frequency across populations of *D. melanogaster*. If abundant precipitation causes NPP to vary spatially and temporally across the range of this species, genotypes better able to find food might prevail in environments with low NPP. Indeed, Williams et al. (2004) found that food shortages in nature may often favor increased mobility in individuals of *D. melanogaster* to find food sources. Specifically, the authors showed evidence that flies become highly active when deprived of food. Yet, some organisms can adjust their foraging behavior to meet environmental changes in food availability (Schoener, 1971). For example, northern pike and white-tailed deer may reduce food intake during winter, when food availability is low (Johnson, 1966; Silver et al., 1969). In the context of the rover-sitter variants, a mechanism known as balancing selection could explain the maintenance of these variants if they are exposed to environmental heterogeneity. Although this mechanism seems plausible in Drosophila, a recent study suggested that strong balancing selection at the *for* gene beginning recently (i.e., following migration out of Africa) is unlikely (Turner et al., 2015). The effect of precipitation as a spatially selective force could not be ruled out though, as the balancing selection test performed by the authors was based on a lineage-specific McDonald-Kreitman model, which does not consider the effect of the environment.

The positive correlations between genetic distance with both geographic and precipitation distances indicate that genetic differentiation between populations is consistent with signatures of IBD and potentially IBE. Accordingly, gene flow may be strong among populations that occur in close proximity and exploit similar environments. Different patterns of gene flow are expected under IBE and IDB models (Sexton et al., 2014). One scenario predicts that with limited dispersal and no selection, genetic drift would cause populations to become more differentiated at greater distances, and that this process should be more pronounced as mean population size decreases (Wright, 1943). Thus, under strict IBD, distance predicts differentiation as a result of dispersal limitation and drift, irrespective of environmental differences. A second scenario predicts gene flow to be strong among similar environments to a greater extent than predicted under IBD. However, adaptation to local environments can disrupt patterns of IBD, and subsequently IBE could arise if maladapted immigrants from different environments are selected against, causing strong barriers to gene flow. A good example of this scenario comes from plants growing on and near mine tailings, where adaptation to different soil types has occurred (Antonovics, 1968). McNeilly and Antonovics (1968) showed that reproductive barriers arose through differences in flower bud development as a result of local adaptation to different soil types. Interestingly, a pattern of isolation by geographical and environmental factors has been reported recently in *D. melanogaster* (Yue et al., 2021). Using microsatellite genotypes, the authors showed an effect of precipitation on patterns of genetic variation across a latitudinal gradient. Similarly, a significant correlation between pairwise *F_st_* and geographic distance was discussed in the context of IBD in a study that examined putatively neutral SNPs, mitochondrial haplotypes, as well as inversion and transposable element insertion polymorphisms (Kapun et al., 2020).

Currently, a number of studies have demonstrated that the foraging behavior of *D. melanogaster* can be strongly influenced by the *for* gene, but none of them have explored the environmental drivers of variation at *for* based on a genotype-environment association study. To my knowledge, this study provides the first evidence that naturally-occurring populations of *D. melanogaster* are genetically differentiated at the *for* gene as a result of environmental selective forces. Importantly, the patterns of genetic differentiation at the *for* gene among populations remained evident when controlling for the genetic variation of the genomic background. Because the samples used in this study consisted of consensus sequences derived from pooled samples (33-40 flies), identifying foraging strains within populations could not be possible (sitters vs rovers). Regardless of this limitation, I showed evidence of an important genetic variation at the *for* gene in this study, but further evidence is required to support a hypothesis of a differential frequency of these strains in nature that can be attributed to balancing selection. Lastly, field studies could disentangle the contributions of selection and environmental structuring to patterns of gene flow. For example, one can conduct an experimental transplant study to understand how dispersal and habitat-specific selection interact to influence populations occupying heterogeneous environments. Although IBE and strong selection should promote local genetic adaptation, it is still unclear how often this is the case in Drosophila.

## Acknowledgements

I thank Drs Michael Angilletta, Jay Taylor, Joaquin Nunez, and John VandenBrooks for providing helpful comments on earlier versions of this manuscript. I also thank the Drosophila Genome Nexus project, the European Drosophila Population Genomics Consortium, and the Real Time Evolution Consortium for providing easy access to the data used in this study.

## Data Accessibility Statement

A fully reproducible workflow of the data analyses, including R scripts and additional supporting material, is available in the following repositories: Github https://dylan-padilla.github.io/for-evo/.

## Conflict of interest

The author has declared no competing interests.

## Author Contributions

Dylan Padilla: Conceptualization, data curation, and formal analysis. Writing – original draft, writing – review and editing. The author agreed to be held accountable for the work performed herein.

## Supplementary material

**Table S1:** Number of seasons surveyed across localities. Abbreviations are as follows: Aus: Austria, Charl = Charlottesville, Denm = Denmark, Esp = Esparto, Fin = Finland, Fran = France, Germ = Germay, PA = Pennsylvania; MI-WI = Michigan and Wisconsin; NY-MA = New York and Massachusetts, Tuol = Tuolume, Ukr = Ukraine.

|  | Aus | Charl | Denm | Esp | Fin | Fran | Germ | MI-WI | NY-MA | PA | Russia | Spain | Tuol | Turkey | Ukr |
| --- | --- | --- | --- | --- | --- | --- | --- | --- | --- | --- | --- | --- | --- | --- | --- |
| fall | 3 | 3 | 4 | 3 | 2 | 4 | 6 | 5 | 3 | 9 | 3 | 4 | 4 | 4 | 14 |
| spring | 3 | 3 | 1 | 2 | 4 | 6 | 6 | 4 | 3 | 8 | 3 | 3 | 1 | 9 | 24 |

## Notes

https://dylan-padilla.github.io/for-evo/

### Competing Interest Statement

The authors have declared no competing interest.

### Summary of Updates

Modifications of figures and text

